# Spatial turnover amplifies with trophic level in hyperdiverse food webs

**DOI:** 10.64898/2026.07.10.732889

**Authors:** Martin Libra, Vojtech Novotny, James B. Whitfield, Scott E. Miller, Ace North, Ondrej Mottl, Philip T. Butterill, Hiroshi Shima, Donald L.J. Quicke, David Wahl, George D. Weiblen, Yves Basset, John Auga, Kenneth Molem, Jan Hrcek

## Abstract

One of the most intuitive ideas in ecology is that diversity at lower trophic levels in food webs provides niches to support diversity at higher trophic levels. This accumulation of diversity can be limited by survival of species in the landscape, but revealing these limits has been challenging. We analyze spatial turnover in a hyperdiverse parasitoid-caterpillar-plant food web across 75,000 km^2^ of continuous lowland rainforest in Papua New Guinea. Species turnover across sites is higher in parasitoids than in their caterpillar hosts. Furthermore, turnover of interactions is also higher in parasitoid-caterpillar than caterpillar-plant networks. Spatial turnover thus amplifies upwards across trophic levels, forcing parasitoids to live closer to spatial persistence limits. Consequently, progressing rainforest fragmentation can especially endanger parasitoids.

## Main text

“Diversity begets diversity” in food webs as the species diversity at one trophic level can promote the diversity of higher trophic levels by providing additional niches (*1, 2*). The co-diversification of plants and herbivorous insects, that led to the emergence of a large part of current global diversity, is an example of this process (*3*). This accumulation of diversity could seemingly continue indefinitely. A potential limitation is that species have to survive in the everyday struggle for existence governed by colonizations and extinctions in local habitat patches (*4*). This may be more difficult for higher trophic levels which are dependent on lower trophic levels. Therefore, comparing spatial turnover between habitat patches among trophic levels can help us understand the underlying ecological processes that limit diversity in food webs.

Species at higher trophic levels experience stronger population instability and greater resource unreliability, which could constrain their diversity (*1*). First, a consumer in a food web can only be present when its resource species are present, limiting the number of habitat patches where it can occur. Second, species from higher trophic levels typically have smaller populations (*5, 6*) and may thus be more prone to local extinctions. Third, consumers can overexploit their resources, increasing the likelihood of local extinction of both interacting species. Species at higher trophic levels, limited by these constraints, are therefore likely to have more patchy occurrence, and some may not be able to sustain viable populations regionally. Increasingly patchy occurrence implies higher difference in community composition among habitat patches. Consequently, we expect species turnover across space to increase from lower to higher trophic levels. At higher trophic levels, spatial turnover may become so pronounced that regional persistence of species becomes increasingly difficult. Very high spatial turnover can therefore signal that the trophic level is approaching the spatial limits of diversity persistence.

Here, we test the expectation of higher spatial turnover at higher trophic level within a hyperdiverse tropical plant–caterpillar–parasitoid food web. We explore spatial turnover across 75,000 km^2^ of highly continuous lowland rainforest in Papua New Guinea, characterized by climatic and floristic uniformity and a lack of pronounced abiotic variation or environmental gradients (Fig. S1, S2, S3). This unique scale of sampling allows us to separate diversity turnover due to interacting trophic levels from the effect of environmental gradients, which themselves strongly influence species turnover (*7*–*10*), as well as from the effects of anthropogenic habitat fragmentation. We investigated the food webs of parasitoids and their caterpillar hosts collected from 25 host plant species at eight sites (Fig. 1A). We focus on the comparison of diversity turnover between caterpillars and their parasitoids, which kill caterpillars by developing inside them (*11*). This comparison is particularly appropriate since caterpillars and their parasitoids have similar body sizes and generation times, unlike, e.g., plants and caterpillars. We reared 27,537 externally feeding caterpillars (Lepidoptera) of 422 species, including 2,875 caterpillars that were parasitized by 304 species of wasp or fly parasitoids (Fig. 1B, Table S1).

**Fig. 1.**
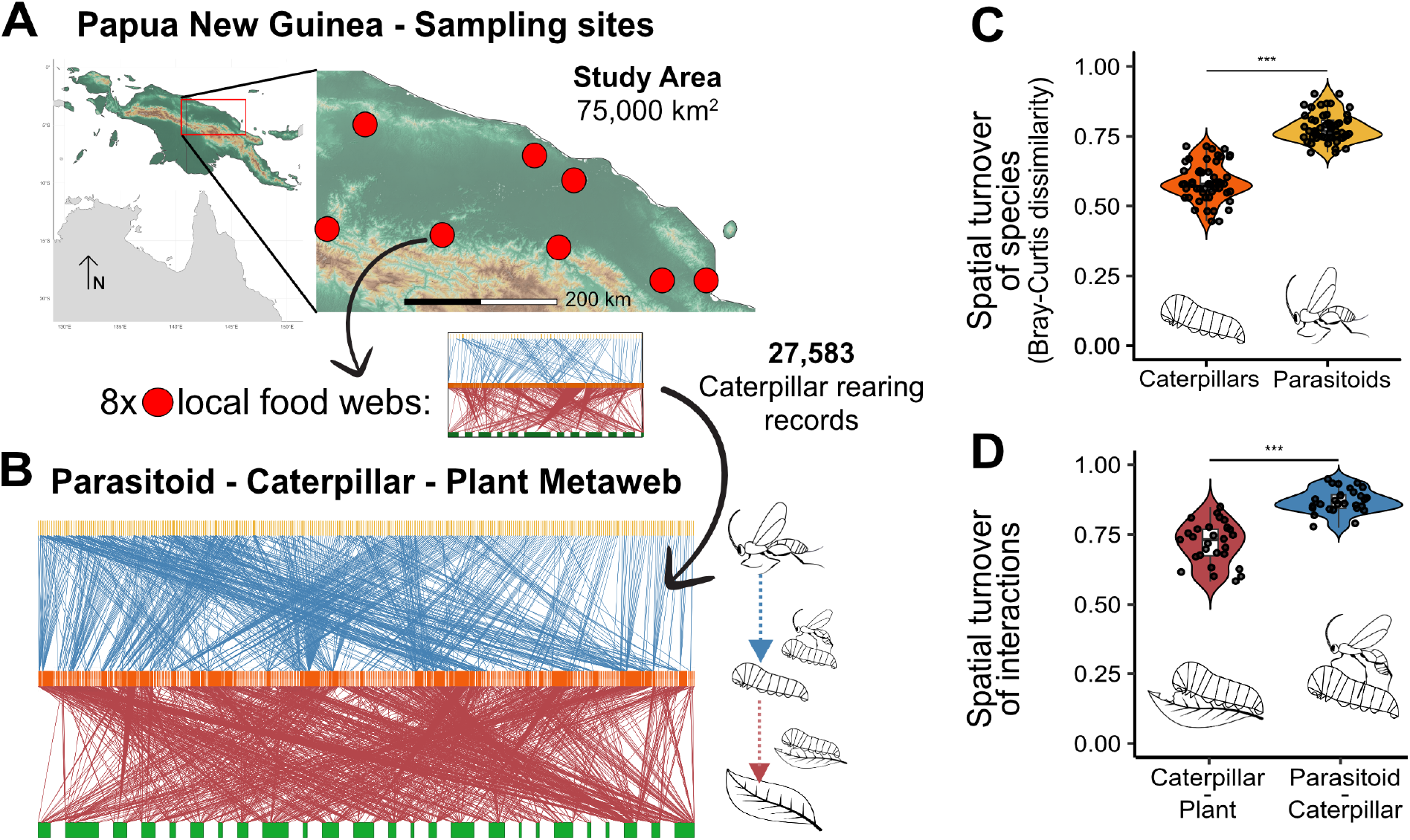
Spatial turnover of caterpillars and parasitoids and their food web interactions across the lowland tropical rainforest of Papua New Guinea. (A) Study area with locations of the eight study sites. (B) Metaweb of all recorded trophic interactions between three trophic levels: plants, caterpillars and parasitoids. The metaweb combines recorded interactions from all eight local food webs. Each bar represents a different species, with width representing sampling effort in plants and abundances in caterpillars and parasitoids. Links represent frequency of recorded interactions between plants and caterpillars (red), or caterpillars and parasitoids (dark blue). (C) Spatial turnover of caterpillar and parasitoid species across sites expressed using Bray–Curtis pairwise dissimilarity of communities. (D) Spatial turnover of food web interactions among sites, shown for Caterpillar–Plant and Parasitoid–Caterpillar interactions. Each point within violin plots (C, D) represents a site-pair dissimilarity (Table S2), ***P < 0.001, Wilcoxon test.

We found that spatial turnover was higher in parasitoids than in their caterpillar hosts across the extensive study area (Fig. 1C). The species turnover in space thus increases with trophic level, consistent with our prediction. High spatial turnover is usually a footprint of high heterogeneity of the abiotic environment (*12*). Yet in our case, the abiotic environment is largely homogeneous. Therefore, heterogeneity in biotic interactions is primarily responsible for high parasitoid spatial turnover.

As comparisons of spatial turnover can be confounded by differences in sample size and proportion of rare species, we conducted a robustness analysis (*13*). The spatial turnover of parasitoids remained higher than that of caterpillars even after downsampling caterpillars to parasitoid sample size (Fig. S4, Table S2). Further, excluding rare parasitoid species or parasitoids from rare caterpillar species or less common host plants had little impact on spatial turnover of parasitoids (Fig. S3, Table S2). The results also remain robust to the choice of spatial turnover measures, which emphasize different aspects of diversity turnover (Fig. S4), indicating a fundamental ecological pattern.

Considering how species interact at individual sites provides additional insight to analysis of species occurrence reported above. We recorded the interactions thanks to the mass rearing we conducted and could therefore reconstruct interaction food webs for each site and compare them (Fig. 1A). Spatial turnover of interactions was significantly higher in the parasitoid-caterpillar food webs than in the caterpillar-plant food webs (Fig. 1D, Fig. 2, Table S4), recapitulating the pattern observed in species occurrence. The pattern of spatial turnover of interactions was again robust to sample size correction (Caterpillar-Plant-subsampled in Fig. 2) and to removal of rare caterpillars and plants (Fig. S5).

**Fig. 2.**
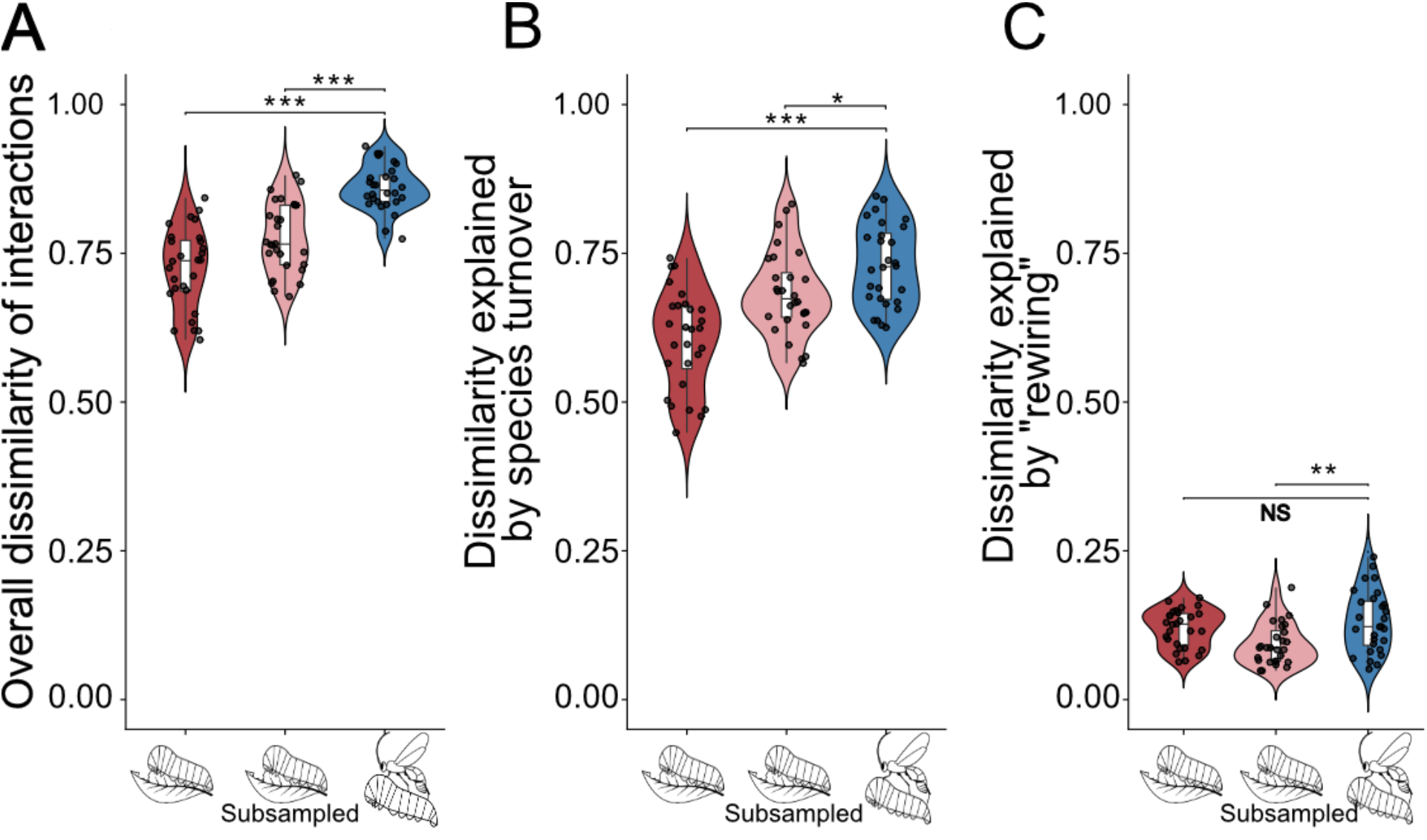
Spatial turnover of food web interactions across the lowland tropical rainforest. A) Overall dissimilarity in interactions between pairs of localities. B) Dissimilarity explained by difference in species turnover between localities. C) Dissimilarity explained by rewiring of interactions to different partners. In addition to Caterpillar-Plant (red), and Parasitoid-Caterpillar (dark blue) interactions we show a Caterpillar-Plant interaction dataset subsampled to the size of the Parasitoid-Caterpillar dataset to remove the potential effect of sample size (Caterpillar-Plant subsampled, pink). Each point within violin plots represents a site-pair dissimilarity (see Methods and (*14*) for dissimilarity computation). In the Caterpillar-Plant subsampled dataset, the points represent the mean value of 1,000 subsamplings of each pairwise dissimilarity. ***P < 0.001, **P < 0.01, *P < 0.05, NS non-significant, Wilcoxon test.

Differences in specialization can contribute to differences in spatial turnover (*13*). Specifically, if parasitoids were more specialized than caterpillars, we would expect higher spatial turnover in parasitoids. We therefore analyzed network-level specialization corrected for sample size and found that the caterpillar–plant food webs were more specialized than parasitoid-caterpillar food webs (H2’ per locality mean ± SD: 0.84 ± 0.04 vs. 0.72 ± 0.10, Wilcoxon test : P = 0.021; Fig. S10). This indicates that specialization does not explain high spatial turnover in parasitoids.

Spatial turnover of interactions can be partitioned into components caused by species turnover and rewiring of interactions to different partners (*14*). In our food webs, spatial turnover of interactions was dominated by species turnover, with low rewiring (Fig. 2BC). This indicates high interaction fidelity across space - when an interaction partner was absent, interactions typically vanished rather than rewired to a new partner. The consequence of this minimal rewiring for parasitoids is more fragmented resources and thus smaller parasitoid populations that are more vulnerable to extinction (*15, 16*).

The low rewiring of parasitoid-caterpillar networks may limit top-down impact on caterpillars from the parasitoids. Hosts can temporarily escape parasitoids by moving to a new location, creating a temporal refuge or “enemy-free space”, as the parasitoids already present there are unlikely to switch to the new host (*17*). We found this enemy-free space to be extensive. On average, 46% of occupied sites represented “enemy-free” locations for a given caterpillar species, lacking any parasitoid species able to attack it (Fig. S8A). Further, a particular caterpillar species was free from attack by a given parasitoid species on average at 67% of the sites it occupied, even though other parasitoids capable of attacking it may have been present (Fig. S8B). As sampling of tropical hyperdiverse food webs is never exhaustive (*18*), very low abundance parasitoids may have gone undetected, but these would have a negligible impact on the hosts. Our data demonstrate that enemy-free space from parasitoids is a pervasive aspect of tropical rainforest herbivore community dynamics.

Yet, when parasitoids are present, their impact can be substantial. On average, parasitoids caused 9.7% (± 12.1 SD) mortality per caterpillar species, but the impact on individual caterpillar species was highly variable among sites and ranged up to 100% mortality (Fig. S9). Total mortality experienced by a caterpillar population at a given site increased with the number of parasitoid species attacking it at that site, with each additional parasitoid species raising the odds of parasitism by 26% (Fig. S11, β = 0.23, P < 0.001). Absence of a parasitoid species from the local community is therefore not compensated by attacks from other parasitoid species and contributes to high spatial variability in parasitoid impact on caterpillars.

The strong spatial turnover in parasitoid communities and high variability in parasitoid impact could be a consequence of temporal dynamics. We therefore analyzed temporal turnover at the Ohu site which we sampled twice. The temporal turnover over three months was higher for parasitoids than for caterpillars, mirroring the results for spatial turnover (Fig. S6A, Table S3). Temporal turnover of interactions was also higher for parasitoid-caterpillar food webs than for caterpillar-plant food webs (Fig. S6B). Caterpillars are an ephemeral resource, susceptible to parasitoids during only 40-60% of the moth/butterfly life cycle (*19*). Hence parasitoids must track suitable hosts across the landscape, possibly resulting in fast dynamics of colonization and extinction.

To better understand how spatial turnover relates to temporal dynamics in food webs, we created a metacommunity model based on stochastic patch occupancy ((*20*); Fig. 3., see Methods). In the model, a number of resource and consumer species inhabit a landscape composed of habitat patches. The resource species are able to colonize nearby patches, whereas consumers are only able to colonize nearby patches if a resource species is present. The model is formulated generally for consumer and resource species and also applies to our more specific case of parasitoids and caterpillars where the local patches represent host plants. As colonization rates are largely unknown for tropical caterpillars (Lepidoptera) and parasitoids, we simulated the metacommunity structure across a wide range of colonization rates of both consumer and resource species. The results of the model simulations show that consumers occupy fewer patches and have a higher spatial turnover than resource species (Fig. 3BC). Indeed, we found near comparable spatial turnover between the two trophic levels only in the unlikely scenario that consumers are consistently much better colonizers than resource species (Fig. 3D). Consumers occupy fewer patches than resource species because the set of patches available to them, at any time, is a subset of those available to resource species. This lower patch occupancy among consumers means that there is less overlap in consumer assemblages between patches, and therefore higher spatial turnover.

**Fig. 3.**
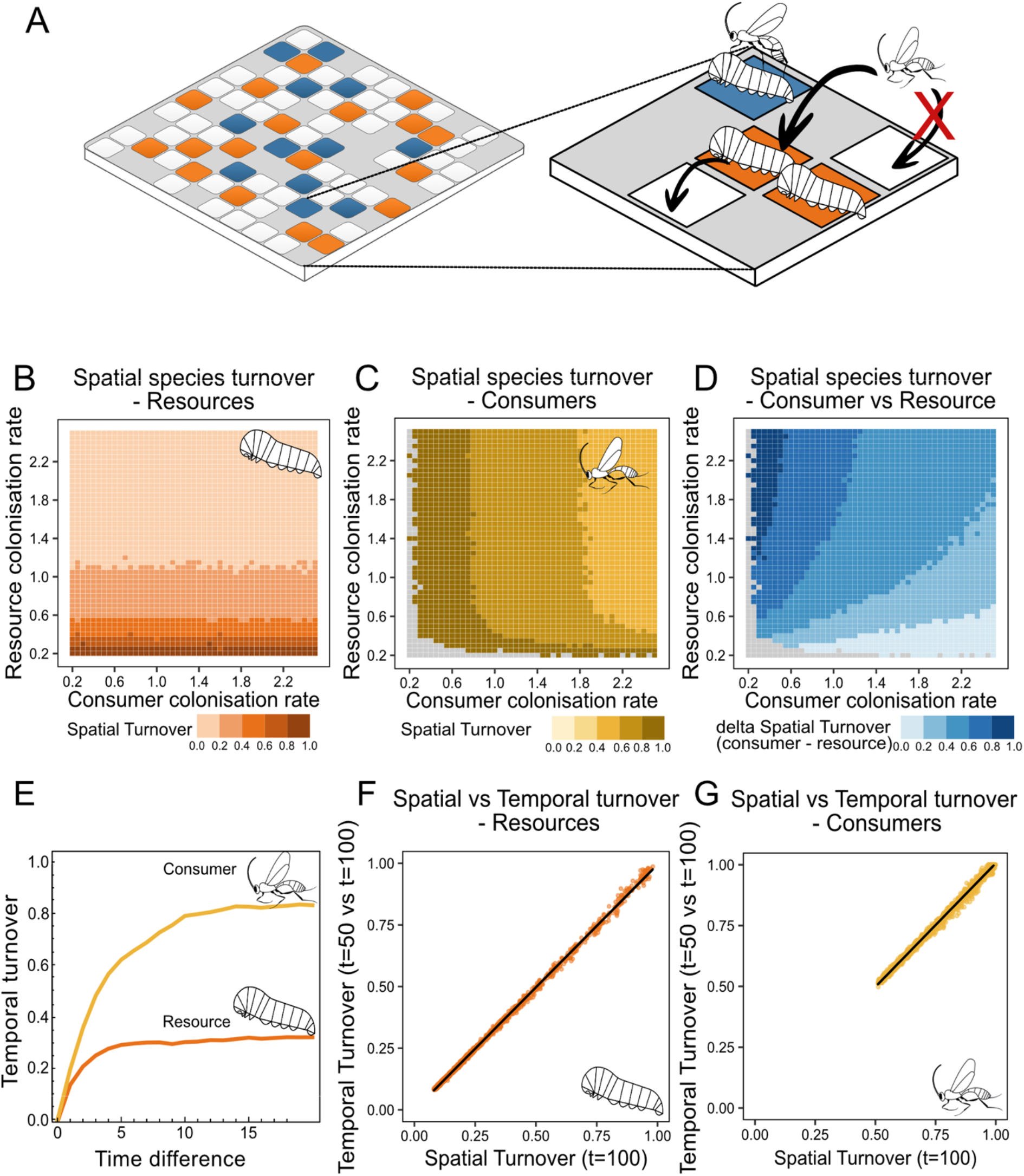
Metacommunity model for consumer-resource interactions. (A) Diagram of the model: 100 patches that resource species can occupy are distributed randomly across a unit square. Consumers can only colonize patches where resource species are present. See Methods for model description and simulation parameters. (B) and (C) show the spatial turnover of resources and consumers (Sørensen index calculated across patches) calculated at the end of the simulation for a range of colonization rates. Grey squares indicate parameter combinations where the resulting community is so sparse that the Sørensen index cannot be computed (there are less than two occupied patches). (D) The difference between spatial turnover in consumers and in resources. (E) Increase of resource and consumer temporal turnover (Sørensen index calculated across time at each patch) with increasing time difference going backward from the end of the simulation. E.g., time difference of 5 represents temporal turnover between timepoints 95 and 100. Temporal turnover is shown for a single example parameter combination of resource and consumer colonization rate of 0.05. (F) and (G) show the correlation between spatial turnover at the end of simulation (timepoint 100) with temporal turnover between timepoint 50 and 100 for resources and consumers, across all colonization rate combinations.

We then explored how consumer and resource communities change in time during the simulations. The difference in community structure increases over time, but more sharply for consumers than for resource species (Fig. 3E). This means that the community turnover is faster for consumers. Eventually, the temporal turnover converges to the level of turnover observed across space (Fig. 3FG). Our model thus provides evidence that a dynamic consumer – resource metacommunity process generates patterns of spatial turnover that amplify with increasing trophic level.

Synthesizing our empirical and theoretical results shows that higher spatial turnover in parasitoids is a parsimonious consequence of modelled metacommunity dynamics. The simple model with a basic assumption that parasitoids occur only where their caterpillars are present readily reproduces the observed pattern. This suggests that amplification of spatial turnover with increasing trophic level may be a widespread phenomenon. The model indicates that parasitoids undergo faster temporal turnover than their caterpillar hosts, and the observed temporal turnover on the Ohu site corroborates this prediction. The higher parasitoid spatial turnover is thus likely a direct consequence of faster extinction–colonization dynamics at the higher trophic level.

Our results are consistent with resource unreliability and population instability constraining the diversity of the higher trophic level, as originally suggested by Whittaker (*1*). Parasitoids are absent from a large proportion of sites occupied by their hosts, revealing challenges they encounter in tracking host populations. The amplification of spatial turnover with increasing trophic level therefore imposes a limit for survival of species at the higher trophic level and explains why the “diversity begets diversity” process cannot continue indefinitely.

The observed amplification of spatial turnover with increasing trophic level can have consequences for adaptation and speciation. The presence or absence of parasitoids at host patches causes large differences in selection pressures experienced by the local host populations. This difference in parasitoid pressure thus creates a geographic mosaic of selection (*21*), even in an environment lacking abiotic gradients. Abrupt changes following parasitoid colonization of a patch create considerable potential for rapid evolution of resistance, potentially leading to eco-evolutionary dynamics distinct from purely ecological host-parasitoid dynamics (*22*). Such micro eco-evolutionary dynamics could be transient or could affect diversification observed at long temporal scales (*23*). We know that half of the common caterpillar species in our study system have spatially structured populations (*24*), but whether this diversification is temporary or indicates progressing speciation is not clear.

The already high spatial turnover of specialized consumers like parasitoids is likely to further increase in response to anthropogenic habitat fragmentation (*25, 26*). Local colonization rates may be reduced, and extinction rates increased to the point when parasitoid metapopulations no longer stay viable. The low rate of interaction rewiring we observed indicates that increasing their host ranges in response to disturbation may not be feasible for parasitoids. Although parasitoids are ecologically important, we know much less about their ecology and conservation status than we know about their hosts on lower trophic levels (*27*). Parasitoids may be one of the more sensitive groups contributing to insect decline (*28*). Our results highlight the urgent need to prioritize studying metapopulation dynamics of higher trophic levels to effectively conserve their biodiversity.

## Acknowledgments

This research was conducted on the traditional lands of indigenous communities in Papua New Guinea. We gratefully acknowledge the contributions of local landowners and community members, whose assistance in field logistics, specimen collection, and processing was indispensable to the success of this study. This research was supported by the National Science Foundation under Grants DEB 9628840, 9707928, 0211591, and 0515678, the Czech Science Foundation (206/09/0115 and 206/08/H044), Czech Academy of Sciences (AA600960712 and AV0Z50070508), Czech Ministry of Education (LC06073, ME916, ME9082, and MSM6007665801), and the Darwin Initiative for the Survival of Species. DNA barcoding was provided by the Centre for Biodiversity Genomics, University of Guelph, with funding from Genome Canada as part of the International Barcode of Life project. ML was supported by ERC grant (669609) and IBERA program 587510/4146. VN was funded by the Praemium Academiae grant, OM by the Charles University Research Centre program (UNCE/24/SCI/006), and by the Institutional Support for Science and Research of the Ministry of Education, Youth and Sports of the Czech Republic. GDW by the David and Lucile Packard Fellowship in Science and Engineering, JH was funded by the European Union (ERC, EcoEvoDiv, 101088709). We thank D. Adamski, V. Becker, J. Brown, K. Darrow, L. Helgen, M. Horak, D. Lees, G. Martin, J.Y. Miller, D. Pollock, M. Rosati, M. Shaffer, N. Silverson, T. Simoes, M.A. Solis, J. Tennent, K. Tuck, and especially J.D. Holloway and J. Rota, and the Natural History Museum (London), for taxonomic assistance with Lepidoptera. We thank C. Godfray, O. Lewis, C. Terry, J. Polechova, A. Buček, and D. Storch for helpful comments on the manuscript.

## References

1. R. H. Whittaker, Evolution and measurement of species diversity. TAXON 21, 213–251 (1972). doi:10.2307/1218190

2. P. A. Hambäck, N. Janz, M. P. Braga, Parasitoid speciation and diversification. Curr. Opin. Insect Sci. 66, 101281 (2024). doi:10.1016/j.cois.2024.101281

3. P. R. Ehrlich, P. H. Raven, Butterflies and plants: a study in coevolution. Evolution 18, 586–608 (1964). doi:10.1111/j.1558-5646.1964.tb01674.x

4. G.-C. Lei, I. Hanski, Metapopulation structure of Cotesia melitaearum, a specialist parasitoid of the butterfly Melitaea cinxia. Oikos 78, 91–100 (1997). doi:10.2307/3545804

5. M. L. Pace, J. J. Cole, S. R. Carpenter, J. F. Kitchell, Trophic cascades revealed in diverse ecosystems. Trends Ecol. Evol. 14, 483–488 (1999). doi:10.1016/S0169-5347(99)01723-1

6. T. Tscharntke, B. A. Hawkins, Multitrophic level interactions: an introduction (Cambridge University Press, Cambridge, 2002).

7. S. C. Maunsell, R. L. Kitching, C. J. Burwell, R. J. Morris, Changes in host–parasitoid food web structure with elevation. J. Anim. Ecol. 84, 353–363 (2015). doi:10.1111/1365-2656.12285

8. G. Peralta, C.M. Frost, R.K. Didham. Plant, herbivore and parasitoid community composition in native Nothofagaceae forests vs. exotic pine plantations. J Appl Ecol. 2018; 55,1265–1275. doi:10.1111/1365-2664.13055

9. M. Seymour, T. Roslin, J. R. deWaard, K. H. J. Perez, M. L. D’Souza, S. Ratnasingham, M. Ashfaq, V. Levesque-Beaudin, G. A. Blagoev, B. Bukowski, P. Cale, D. Crosbie, T. Decaëns, S. L. deWaard, T. Ekrem, H. O. El-Ansary, F. Evouna Ondo, D. Fraser, M. F. Geiger, M. Hajibabaei, W. Hallwachs, P. E. Hanisch, A. Hausmann, M. Heath, I. D. Hogg, D. H. Janzen, M. Kinnaird, J. R. Kohn, M. Larrivée, D. C. Lees, V. León-Règagnon, M. Liddell, D. A. Lijtmaer, T. Lipinskaya, S. A. Locke, R. Manjunath, D. J. Martins, M. B. Martins, S. Mazumdar, J. T. A. McKeown, K. Anderson-Teixeria, S. E. Miller, M. A. Milton, R. Miskie, J. Morinière, M. Mutanen, S. Naik, B. Nichols, F. A. Noguera, V. Novotny, L. Penev, M. Pentinsaari, J. Quinn, L. Ramsay, R. Rochefort, S. Schmidt, M. A. Smith, C. N. Sobel, P. Somervuo, J. E. Sones, H. S. Staude, B. St. Jaques, E. Stur, A. C. Telfer, P. L. Tubaro, T. J. Wardlaw, R. Worcester, Z. Yang, M. R. Young, T. Zemlak, E. V. Zakharov, B. Zlotnick, O. Ovaskainen, P. D. N. Hebert, Global arthropod beta-diversity is spatially and temporally structured by latitude. Commun. Biol. 7, 552 (2024). doi:10.1038/s42003-024-06199-1

10. J. O. Stireman III, The evolution of generalization? Parasitoid flies and the perils of inferring host range evolution from phylogenies. J. Evol. Biol. 18, 325–336 (2005). doi:10.1111/j.1420-9101.2004.00850.x

11. M. A. Condon, S. J. Scheffer, M. L. Lewis, R. Wharton, D. C. Adams, A. A. Forbes, Lethal interactions between parasites and prey increase niche diversity in a tropical community. Science 343, 1240–1244 (2014). doi:10.1126/science.1245007

12. V. Fontana, E. Guariento, A. Hilpold, G. Niedrist, M. Steinwandter, D. Spitale, J. Nascimbene, U. Tappeiner, J. Seeber, Species richness and beta diversity patterns of multiple taxa along an elevational gradient in pastured grasslands in the European Alps. Sci. Rep. 10, 12516 (2020). doi:10.1038/s41598-020-69569-9

13. L. Pereira Martins, A. Matos Medina, T. M. Lewinsohn, M. Almeida-Neto, Trophic level and host specialisation affect beta-diversity in plant–herbivore–parasitoid assemblages. Insect Conserv. Divers. 12, 404–413 (2019). doi:10.1111/icad.12353

14. T. Poisot, E. Canard, D. Mouillot, N. Mouquet, D. Gravel, The dissimilarity of species interaction networks. Ecol. Lett. 15, 1353–1361 (2012). doi:10.1111/ele.12002

15. C. T. Jeffs, O. T. Lewis, Effects of climate warming on host–parasitoid interactions. Ecol. Entomol. 38, 209–218 (2013). doi:10.1111/een.12026

16. S. van Nouhuys, Effects of habitat fragmentation at different trophic levels in insect communities. Ann. Zool. Fenn. 42, 433–447 (2005).

17. M. J. Jeffries, J. H. Lawton, Enemy free space and the structure of ecological communities. Biol. J. Linn. Soc. 23, 269–286 (1984). doi:10.1111/j.1095-8312.1984.tb00145.x

18. V. Novotny, S. E. Miller, L. Baje, S. Balagawi, Y. Basset, L. Cizek, K. J. Craft, F. Dem, R. A. I. Drew, J. Hulcr, J. Leps, O. T. Lewis, R. Pokon, A. J. A. Stewart, G. Allan Samuelson, G. D. Weiblen, Guild-specific patterns of species richness and host specialization in plant–herbivore food webs from a tropical forest. J. Anim. Ecol. 79, 1193–1203 (2010). doi:10.1111/j.1365-2656.2010.01728.x

19. D. L. Wagner, Caterpillars of Eastern North America (Princeton University Press, NJ, 2005).

20. O. Ovaskainen, I. Hanski. From individual behavior to metapopulation dynamics: unifying the patchy population and classic metapopulation models. Am Nat. 164, 364–77 (2004). doi: 10.1086/423151

21. J. N. Thompson, The Geographic Mosaic of Coevolution (University of Chicago Press, Chicago, 2005)

22. T. Hiltunen, L. Becks, Consumer co-evolution as an important component of the eco-evolutionary feedback. Nat. Commun. 5, 5226 (2014). doi:10.1038/ncomms6226

23. E. A. Fronhofer, D. Corenblit, J. N. Deshpande, L. Govaert, P. Huneman, F. Viard, P. Jarne, S. Puijalon, Eco-evolution from deep time to contemporary dynamics: The role of timescales and rate modulators. Ecol. Lett. 26, S91– S108 (2023). doi:10.1111/ele.14222

24. K. J. Craft, S. U. Pauls, K. Darrow, S. E. Miller, P. D. N. Hebert, L. E. Helgen, V. Novotny, G. D. Weiblen, Population genetics of ecological communities with DNA barcodes: An example from New Guinea Lepidoptera. Proc. Natl. Acad. Sci. 107, 5041–5046 (2010). doi:10.1073/pnas.0913084107

25. S. P. Hubbell, F. He, R. Condit, L. Borda-de-Água, J. Kellner, H. ter Steege, How many tree species are there in the Amazon and how many of them will go extinct? Proc. Natl. Acad. Sci. 105, 11498–11504 (2008). doi:10.1073/pnas.0801915105

26. H. ter Steege, N. C. A. Pitman, I. L. do Amaral, L. de Souza Coelho, F. D. de Almeida Matos, D. de Andrade Lima Filho, R. P. Salomão, F. Wittmann, C. V. Castilho, J. E. Guevara, M. de J. Veiga Carim, O. L. Phillips, W. E. Magnusson, D. Sabatier, J. D. C. Revilla, J.-F. Molino, M. V. Irume, M. P. Martins, J. R. da Silva Guimarães, J. F. Ramos, O. S. Bánki, M. T. F. Piedade, D. Cárdenas López, D. de J. Rodrigues, L. O. Demarchi, J. Schöngart, E. J. Almeida, L. F. Barbosa, L. Cavalheiro, M. C. V. dos Santos, B. G. Luize, E. M. M. de Leão Novo, P. N. Vargas, T. S. F. Silva, E. M. Venticinque, A. G. Manzatto, N. F. C. Reis, J. Terborgh, K. R. Casula, E. N. Honorio Coronado, A. Monteagudo Mendoza, J. C. Montero, F. R. C. Costa, T. R. Feldpausch, A. C. Quaresma, N. Castaño Arboleda, C. E. Zartman, T. J. Killeen, B. S. Marimon, B. H. Marimon-Junior, R. Vasquez, B. Mostacedo, R. L. Assis, C. Baraloto, D. D. do Amaral, J. Engel, P. Petronelli, H. Castellanos, M. B. de Medeiros, M. F. Simon, A. Andrade, J. L. Camargo, W. F. Laurance, S. G. W. Laurance, L. Maniguaje Rincón, J. Schietti, T. R. Sousa, E. de Sousa Farias, M. A. Lopes, J. L. L. Magalhães, H. E. M. Nascimento, H. L. de Queiroz, G. A. Aymard C. R. Brienen, P. R. Stevenson, A. Araujo-Murakami, T. R. Baker, B. B. L. Cintra, Y. O. Feitosa, H. F. Mogollón, J. F. Duivenvoorden, C. A. Peres, M. R. Silman, L. V. Ferreira, J. R. Lozada, J. A. Comiskey, F. C. Draper, J. J. de Toledo, G. Damasco, R. García-Villacorta Lopes, A. Vicentini, F. Cornejo Valverde, A. Alonso, L. Arroyo, F. Dallmeier, V. H. F. Gomes, E. M. Jimenez, D. Neill, M. C. Peñuela Mora, J. C. Noronha, D. P. P. de Aguiar, F. R. Barbosa, Y. K. Bredin, R. de Sá Carpanedo F. A. Carvalho, F. C. de Souza, K. J. Feeley, R. Gribel, T. Haugaasen, J. E. Hawes, M. P. Pansonato, M. Ríos Paredes, J. Barlow, E. Berenguer, I. B. da Silva, M. J. Ferreira, J. Ferreira, P. V. A. Fine, M. C. Guedes, C. Levis, J. C. Licona, E. Villa Zegarra, V. A. Vos, C. Cerón, F. M. Durgante, É. Fonty, T. W. Henkel, J. E. Householder, I. Huamantupa-Chuquimaco, E. Pos, M. Silveira, J. Stropp, R. Thomas, D. Daly, K. G. Dexter, W. Milliken, G. P. Molina, T. Pennington, I. C. G. Vieira, B. Weiss Albuquerque, W. Campelo, A. Fuentes, B. Klitgaard, J. L. M. Pena, J. S. Tello, Vriesendorp, J. Chave, A. Di Fiore, R. R. Hilário, L. de Oliveira Pereira, J. F. Phillips, G. Rivas-Torres, T. R. van Andel, P. von Hildebrand, W. Balee, E. M. Barbosa, L. C. de Matos Bonates, H. P. Dávila Doza, R. Zárate Gómez, T. Gonzales, G. P. Gallardo Gonzales, B. Hoffman, A. B. Junqueira, Y. Malhi, I. P. de Andrade Miranda, L. F. M. Pinto, A. Prieto, A. Rudas, A. R. Ruschel, N. Silva, C. I. A. Vela, E. L. Zent, S. Zent, A. Cano, Y. A. Carrero Márquez, D. F. Correa, J. B. P. Costa, B. M. Flores, D. Galbraith, M. Holmgren, M. Kalamandeen, G. Lobo, L. Torres Montenegro, M. T. Nascimento, A. A. Oliveira, M. M. Pombo, H. Ramirez-Angulo, M. Rocha, V. V. Scudeller, R. Sierra, M. Tirado, M. N. Umaña, G. van der Heijden, E. Vilanova Torre, M. A. A. Reategui, C. Baider, H. Balslev, S. Cárdenas, L. F. Casas, M. J. Endara, W. Farfan-Rios, C. Ferreira, R. Linares-Palomino, C. Mendoza, I. Mesones, G. A. Parada, A. Torres-Lezama, L. E. Urrego Giraldo, D. Villarroel, R. Zagt, M. N. Alexiades, E. A. de Oliveira, K. Garcia-Cabrera, L. Hernandez, W. P. Cuenca, S. Pansini, D. Pauletto, F. Ramirez Arevalo, A. F. Sampaio, E. H. Valderrama Sandoval, L. V. Gamarra, A. Levesley, G. Pickavance, K. Melgaço, Mapping density, diversity and species-richness of the Amazon tree flora. Commun. Biol. 6, 1130 (2023). doi:10.1038/s42003-023-05514-6

27. K. D. Lafferty, S. Allesina, M. Arim, C. J. Briggs, G. De Leo, A. P. Dobson, J. A. Dunne, P. T. J. Johnson, A. M. Kuris, D. J. Marcogliese, N. D. Martinez, J. Memmott, P. A. Marquet, J. P. McLaughlin, E. A. Mordecai, M. Pascual, R. Poulin, D. W. Thieltges, Parasites in food webs: the ultimate missing links. Ecol. Lett. 11, 533–546 (2008). doi:10.1111/j.1461-0248.2008.01174.x

28. J. A. Harvey, K. Tougeron, R. Gols, R. Heinen, M. Abarca, P. K. Abram, Y. Basset, M. Berg, C. Boggs, J. Brodeur, P. Cardoso, J. G. de Boer, G. R. De Snoo, C. Deacon, J. E. Dell, N. Desneux, M. E. Dillon, G. A. Duffy, L. A. Dyer, J. Ellers, A. Espíndola, J. Fordyce, M. L. Forister, C. Fukushima, M. J. G. Gage, C. García-Robledo, C. Gely, M. Gobbi, C. Hallmann, T. Hance, J. Harte, A. Hochkirch, C. Hof, A. A. Hoffmann, J. G. Kingsolver, G. P. A. Lamarre, W. F. Laurance, B. Lavandero, S. R. Leather, P. Lehmann, C. Le Lann, M. M. López-Uribe, C.-S. Ma, G. Ma, J. Moiroux, L. Monticelli, C. Nice, P. J. Ode, S. Pincebourde, W. J. Ripple, M. Rowe, M. J. Samways, A. Sentis, A. A. Shah, N. Stork, J. S. Terblanche, M. P. Thakur, M. B. Thomas, J. M. Tylianakis, J. Van Baaren, M. Van de Pol, W. H. Van der Putten, H. Van Dyck, W. C. E. P. Verberk, D. L. Wagner, W. W. Weisser, W. C. Wetzel, H. A. Woods, K. A. G. Wyckhuys, S. L. Chown, Scientists’ warning on climate change and insects. Ecol. Monogr. 93, e1553 (2023). doi: 10.1002/ecm.1553

29. V. Novotny, S. E. Miller, J. Hulcr, R. A. I. Drew, Y. Basset, M. Janda, G. P. Setliff, K. Darrow, A. J. A. Stewart, J. Auga, B. Isua, K. Molem, M. Manumbor, E. Tamtiai, M. Mogia, G. D. Weiblen, Low beta diversity of herbivorous insects in tropical forests. Nature 448, 692–695 (2007). doi:10.1038/nature06021

30. J. Hrcek, S. E. Miller, J. B. Whitfield, H. Shima, V. Novotny, Parasitism rate, parasitoid community composition and host specificity on exposed and semi-concealed caterpillars from a tropical rainforest. Oecologia 173, 521–532 (2013). doi:10.1007/s00442-013-2619-6

31. R Core Team, R: A language and environment for statistical computing (R Foundation for Statistical Computing, Vienna, 2021); https://www.R-project.org/.

32. A. Baselga, C. D. L. Orme, betapart: an R package for the study of beta diversity. Methods Ecol. Evol. 3, 808–812 (2012). doi:10.1111/j.2041-210X.2012.00224.x

33. A. Chao, R. L. Chazdon, R. K. Colwell, T.-J. Shen, A new statistical approach for assessing similarity of species composition with incidence and abundance data. Ecol. Lett. 8, 148–159 (2005). doi:10.1111/j.1461-0248.2004.00707.x

34. A. S. Melo, CommEcol: community ecology analyses, R package version 1.7.0 (2013); https://CRAN.R-project.org/package=CommEcol.

35. T. J. Sørensen, A method of establishing groups of equal amplitude in plant sociology based on similarity of species content (E. Munksgaard, Copenhagen, 1948).

36. C. Dormann, J. Fründ, N. Blüthgen, B. Gruber, Indices, graphs and null models: analyzing bipartite ecological networks. Open Ecol. J. 2, 7–24 (2009). doi:10.2174/1874213000902010007

37. A. Baselga, Partitioning the turnover and nestedness components of beta diversity. Global Ecol. Biogeogr. 19, 134–143 (2010). doi:10.1111/j.1466-8238.2009.00490.x

38. D. Bates, M. Mächler, B. Bolker, S. Walker, Fitting linear mixed-effects models using lme4. J. Stat. Softw. 67, 1–48 (2015). doi:10.18637/jss.v067.i01

39. B. Bolker, D. Robinson, broom.mixed: tidying methods for mixed models, R package version 0.2.9.4 (2022); https://CRAN.R-project.org/package=broom.mixed

40. J. T. Abatzoglou, S. Z. Dobrowski, S. A. Parks, K. C. Hegewisch, TerraClimate, a high-resolution global dataset of monthly climate and climatic water balance from 1958–2015. Sci. Data 5, 170191 (2018). doi:10.1038/sdata.2017.191

41. Oksanen J, Simpson G, Blanchet F, Kindt R, Legendre P, Minchin P, O’Hara R, Solymos P, Stevens M, Szoecs E, Wagner H, Barbour M, Bedward M, Bolker B, Borcard D, Borman T, Carvalho G, Chirico M, De Caceres M, Durand S, Evangelista H, FitzJohn R, Friendly M, Furneaux B, Hannigan G, Hill M, Lahti L, Martino C, McGlinn D, Ouellette M, Ribeiro Cunha E, Smith T, Stier A, Ter Braak C, Weedon J (2026). vegan: Community Ecology Package. https://CRAN.R-project.org/package=vegan >

42. D. Gravel, E. Canard, F. Guichard, N. Mouquet, Persistence increases with diversity and connectance in trophic metacommunities. PLoS One 6, e19374 (2011). doi:10.1371/journal.pone.0019374

